# Ecological and genetic trade-offs drive *Arabidopsis thaliana* range expansion in Europe

**DOI:** 10.1101/2022.11.29.518298

**Authors:** Cristina C. Bastias, Aurélien Estarague, Denis Vile, Cheng-Ruei Lee, Moises Exposito-Alonso, Cyrille Violle, François Vasseur

**Author notes:** Corresponding author: Cristina C. Bastias., CEFE, Univ Montpellier, CNRS, EPHE, IRD, Montpellier, France. Área de Ecología, Facultad de Ciencias, Universidad de Córdoba, Campus de Rabanales, 14071 Córdoba, Spain. These authors equally contributed to this work (Cristina C. Bastias, Aurélien Estarague). These authors share the senior authorship (Cyrille Violle, François Vasseur).

## Abstract

How trade-offs between traits constrain adaptation to contrasted environments is critical to understand the distribution range of a given species. In *Arabidopsis thaliana*, genetic analyses recently revealed that a group of genotypes successfully recolonized Europe from its center after the last glaciation, outcompeting older lineages and leaving them only at the distribution margins, where environmental conditions are more stressing. However, whether trade-offs between traits related to dispersal, competition, and stress tolerance explain the success and persistence of different lineages across the species geographic range remains an open question. Here, we compared the genetic and phenotypic differentiation between 72 ecotypes originating from three geographical groups in Europe (North, South and Center). We measured key traits related to fecundity, dispersal ability, competition tolerance, and stress tolerance, and used genomic data to infer the effect of selection on these traits. We showed that a trade-off between plant fecundity and seed mass constrains the diversification of *A. thaliana* in Europe. In particular, the success of the cosmopolitan genotypes that recolonized Europe can be explained by their higher dispersal ability at the expense of their competitive ability and stress tolerance. Inversely, peripheral ecotypes exhibited the opposite trait syndrome: high competition and stress tolerance but low dispersal ability. Moreover, peripheral genotypes tend to differentiate from central ones at genes involved in dispersal and competitive traits such as seed mass. Combining ecological and genomic approaches, our study demonstrated the role of key ecological trade-offs as evolutionary drivers of the distribution of plant populations along a geographic gradient.

**Significance:** Across geographic gradients, differential adaptive phenotypes among populations can reduce the risk of local extinctions and favor niche dynamics. However, a phenotypic advantage often comes at a cost. For instance, the competition-colonization trade-off is proposed as an important driver of plant interspecific diversity, but its role for local adaptation at the intraspecific level is still unclear. Using the broadly-distributed species *Arabidopsis thaliana*, we evaluate how ecological trade-offs have shaped the demography and evolution of central and peripheral populations in its native geographic gradient. Our study demonstrates that the competition-colonization trade-off is responsible of the spatially-structured phenotypic variation of *A. thaliana* across its geographical range. Our study highlights seed mass as a key trait for plant adaptation across environmental conditions.

## INTRODUCTION

Ecophysiological considerations suggest that plants cannot simultaneously optimize all major functional traits, resulting in ecological trade-offs between key plant’ functions as dispersal, colonization, competition and stress tolerance^1–3^. Such trade-offs impose limits on plant phenotypic adaptation, which is at the origin of the local extinction processes of numerous plant populations in the face of environmental hazards^4–6^. Trade-offs are also responsible for the contractions or expansions of the species distribution area at the medium and long term^3^. Numerous theoretical expectations about population’s adaptive differentiation across species distribution range have been proposed in ecology and evolution ^4–8^, but empirical evidence remains scarce.

The *competition-colonization trade-off* ^7–9^ relies on the idea that resource limitation imposes plants to face a critical dilemma: producing many small seeds or few big seeds. The first strategy (producing many small seeds) is expected to optimize fecundity, dispersal, and colonization ability^10^, while the second strategy (producing few big seeds) is expected to optimize seedling recruitment and survival and produce more superior competitors^8,11–13^. Beyond seed mass and seed number, this trade-off is expected to be associated with many other traits related to competitive, colonization and dispersal abilities^11,14^, in particular with reproductive plant height^15,16^. Moreover, traits that influence competitive ability, such as plant growth and seed mass, are also expected to determine in large extent plant response to biotic and abiotic stress, thus generating a corollary trade-off referred to as the *colonization-stress tolerance trade-off*^17,18^. Surprisingly, empirical evidence of the competition-colonization trade-off (as well as the colonization-stress tolerance trade-off) remains ambiguous, and mostly limited to large comparative studies across diverse plant species^7,14,19^. Despite their potential role for plant evolution, these trade-offs have been poorly investigated at the intraspecific level. More specifically, if and how trade-offs shape the demography and evolution of plant populations across the geographical range is still unclear.

Thanks to the contribution of modern genomics and genetic analyses at the intraspecific level, evolutionary biologists are now able to trace the demographic history of species. For example, after the last glaciation, many species recolonized Europe from the south via climate-refugee populations^20,21^. Some species have also undergone secondary recolonization of Europe, benefiting from “supercolonizing” variants that quickly spread throughout a large part of the distribution range^22,23^. In such cases, older natural lineages, such as climate-refugee populations, often went extinct and replaced by the cosmopolitan variants. However, in some cases, climate-refugee populations (called ‘relicts’) can persist and cohabit locally with the cosmopolitan (nonrelict) variants and even potentially admix and share adaptive alleles that confer a phenotypic advantage under specific environmental conditions. This has been typically observed in the model plant *Arabidopsis thaliana*^24,25^, which benefits from an unprecedented characterization of its evolutionary history and geographical distribution^26–28^. Recent genomic analyses of *A. thaliana* diversity notably revealed that the vast majority of natural lines across Europe (cosmopolitan groups hereafter) share a common genetic legacy from a secondary colonization from central Europe starting around 10,000 years ago (Fig. 1)^24,25^. Relicts, i.e. highly-diverged groups coming from the Ice Age climate-refugees, have only persisted until now in the southern European areas (most of them in the Iberian Peninsula)^24,25^. Nonetheless, genetic analyses of the 1,001 Genomes dataset^29^ further indicated that the cosmopolitan accessions occupying the North of Scandinavia are genetically closer to relict lines than central European populations in genes linked to drought tolerance^24,30^. This has been explained by the presence of relicts in the northernmost European areas before the last Ice Age^24^. In addition, this suggests that similar selection forces may operate at the opposite edges of the species range, although we do not know the phenotypic traits and the mechanisms involved.

**Fig. 1.**
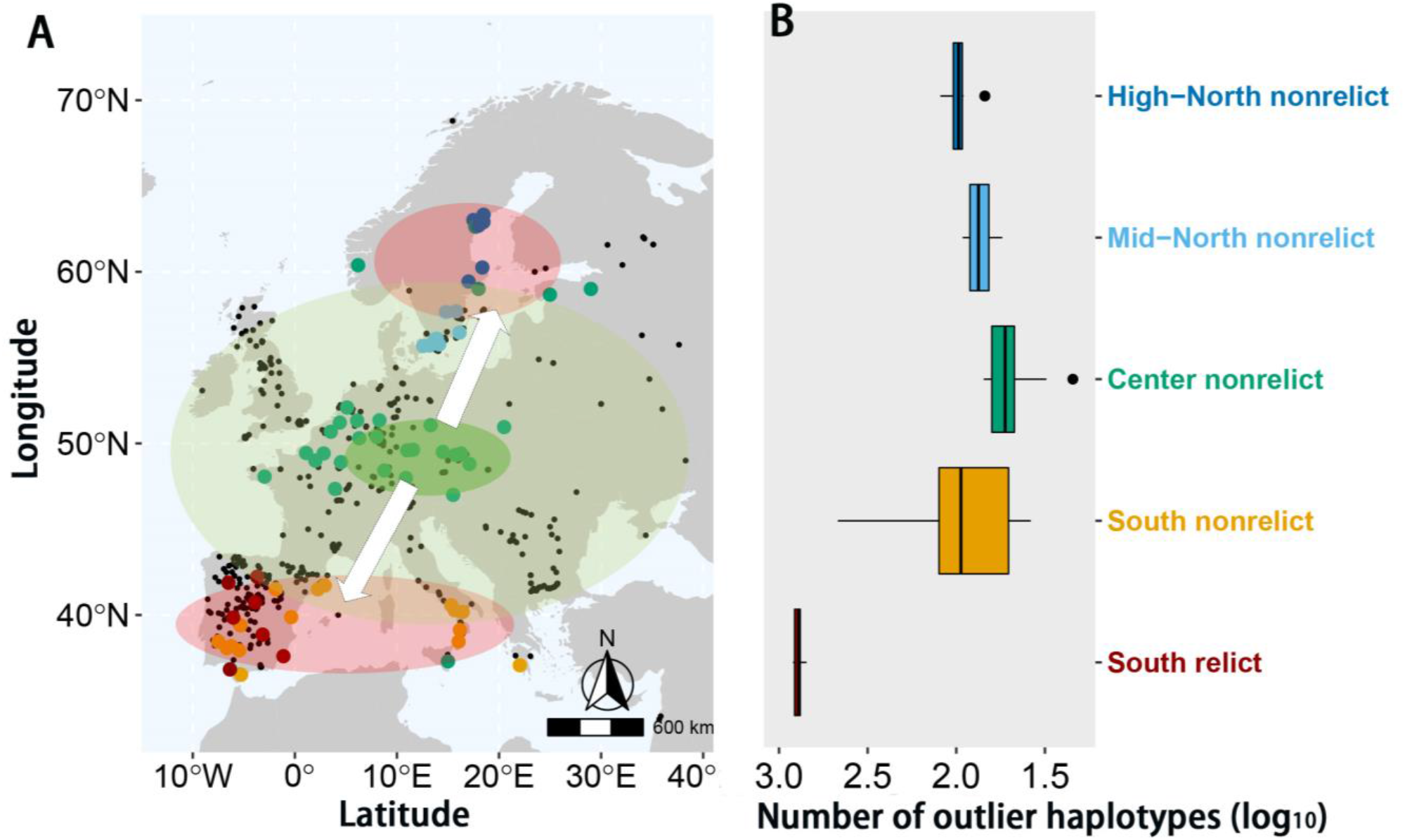
Demographic history of *A. thaliana* in Europe. (A) Lee and collaborators ^24^ suggested an evolutionary scenario based on two expansion waves. First, populations from different glacial refugia expanded northwards reaching scandinavian areas at the end of the last Ice Age in Europe. Afterwards, a second wave of expansion initiated from a nonrelict clade set up at central European areas (dark green circle) and quickly spread out across Europe (light green circle), mainly erasing any trace of those first colonizers (relict groups) at mid latitudes. Later, genetic studies also evidenced that nonrelict accessions occupying the northernmost area in Europe share alleles with relict accessions from southern areas (both represented with red circles). Black points represent natural populations for *A. thaliana* within 1,001 Genomes project in its European distribution (http://1001genomes.org/). Points in color represent the 72 *A. thaliana* selected accessions categorized per geographical group and its genetic basis. (B) Number of outlier haplotypes (log_10_), i.e. number of those alleles highly diverged from all other alleles across the species and make accessions genetically distant from all others^24^, for each *A. thaliana* natural accession categorized in each geographical group.

Genome-wide association studies (GWAS)^31^ enable us to examine the genetic determinism of trait variability across accessions and to test whether a trait is under selection or not. For instance, the *Q*_*ST*_-*F*_*ST*_ approach aims at comparing the degree of phenotypic differentiation (*i*.*e. Q*_*ST*_) to the degree of genetic differentiation at neutral markers (*i*.*e. F*_*ST*_) between populations^32,33^. A *Q*_*ST*_ / *F*_*ST*_ ratio of 1 suggests that the studied trait does not differ between populations more than what we can expect under the effect of genetic drift only. A *Q*_*ST*_ / *F*_*ST*_ > 1 suggests the effect of directional selection on the trait, while *Q*_*ST*_ / *F*_*ST*_ < 1 suggests the effect of stabilizing or uniform selection^33^. For practical reasons, *Q*_*ST*_ has been often measured at the phenotypic level as the additive genetic variation on a trait between populations. However, the large sequencing effort of *A. thaliana* gives the opportunity to identify the single nucleotide polymorphisms (SNPs) involved in trait variation^29^, and then to calculate *Q*_*ST*_ at the genomic level, *i*.*e*. the *F*_*ST*_ of SNPs that impacts on the trait of interest (*F*_*STQ*_ *sensu* ref.^34^).

Here, we hypothesized that the success of the central cosmopolitan (nonrelict) groups in *A. thaliana* was due to their superior colonization ability (*e*.*g*., tall, highly-fecund, and small-seeded accessions), which according to the competition-colonization trade-off should be detrimental to their ability to compete and tolerate stress. By contrast, relict accessions and those ones genetically close to them (*e*.*g*., nonrelicts from North Scandinavia) are expected to exhibit a high competition ability (*e*.*g*., big-seeded accessions) and environmental stress tolerance, which could explain why they persist at the opposite *A. thaliana* range edges. To test these hypotheses, we performed a common garden experiment in greenhouse using 72 natural accessions of *A. thaliana* differing in genetic and geographical distances along its native European distribution range, in order (i) to assess the spatial structure of phenotypic variation in key traits related to dispersal, colonization, competition and stress-response abilities along *A. thaliana* distribution range, (ii) to test whether ecological trade-offs hold at the intraspecific level, and (iii) to determine the genetic and evolutionary bases underlying this phenotypic distribution.

## RESULTS

### Extensive intraspecific trait variability across European distribution range for *A. thaliana*

The 72 natural accessions of *A. thaliana* were equally selected in three European geographical groups. Within each geographical group, accessions were classified as relicts or nonrelicts according to recent genomic-based studies^24,29^. These are ‘Center nonrelict’ group from Central Europe, ‘South relict’ and ‘South nonrelict’ from the southern European areas and ‘Mid-North nonrelict’ and ‘High-North nonrelict’ from the northern European areas (Fig. 1A, Table S1). Using previously published data from Lee and colleagues^24^, we detected that relict accessions contain a high outlier haplotype numbers, i.e. alleles less frequent across whole species genome and make accessions genetically distant from all others^24^ (Fig. 1B). However, the nonrelict groups that are closer to the species distribution edges also exhibited more outlier haplotypes than central nonrelicts. Thus, the geographical distribution of *A. thaliana* is associated with an increase in outlier haplotypes in the genomic background toward the edges (Fig. 1B).

A total of 24 individuals per accession were grown in greenhouse under control conditions (one single plant per pot and well-watering), intraspecific competition (one focal plant surrounded by 4 neighbors of the same accession and well-watering) and under water stress conditions (one single plant with withholding watering for a period of 10-11 days). For each individual, we measured its fecundity (total fruit number x average fruit size), seed mass (averaged from 10-30 seeds) and maximum plant height. The responses to competition and water stress were assessed in terms of fecundity through reaction norms *i*.*e*. the difference in individual values of fecundity in competition or water-stress, and the mean fecundity of the same accession in control conditions. All traits substantially varied across accessions (Fig. S2) and a relevant number (7 out of 10) of significant pairwise correlations were found between our studied traits (Fig. S3).

We specifically found that, across all accessions in control conditions, fecundity was negatively correlated with seed mass (Pearson’s correlation coefficient (*r*) = -0.50, *P* < 0.001; n = 71; Fig. 2 and Fig. S3) and positively correlated with plant height (*r* = 0.51, *P* < 0.001; n = 71; Fig. S3). Fecundity was negatively correlated with intraspecific competition tolerance (*r* = -0.978, *P* < 0.001; n = 71; Fig. S3) and water stress tolerance (*r* = -0.54, *P* < 0.001; n = 71; Fig. S3). Seed mass and competition tolerance were positively correlated (*r* = 0.45, *P <* 0.001; n = 71; Fig. S3). In contrast, seed mass did not correlate with water stress tolerance (*r =* 0.12, *P* = 0.30; n = 71; Fig. S3). Neither plant height nor seed mass was correlated among them or showed correlation with the response to water stress (Fig. S3).

**Fig. 2.**
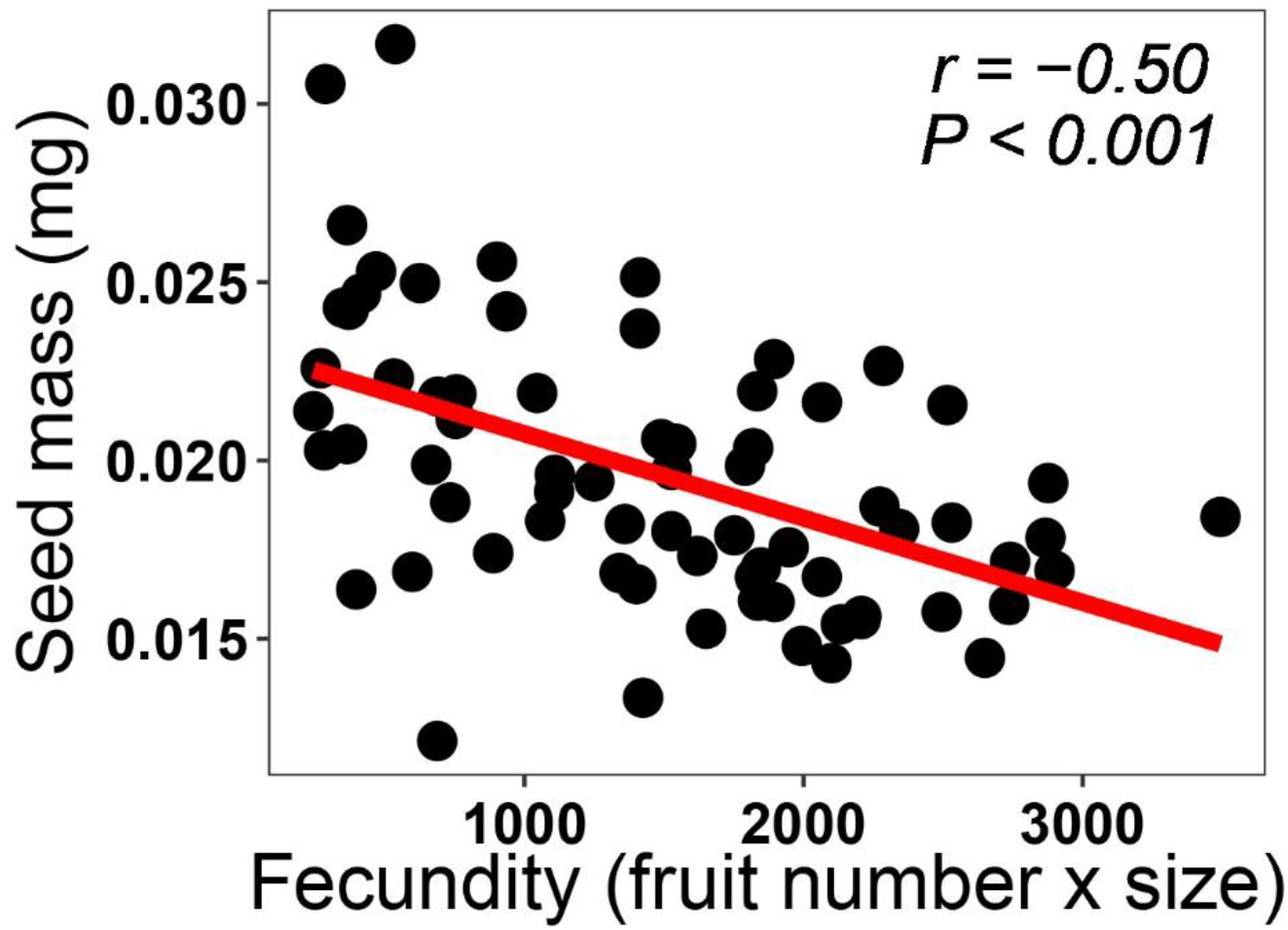
Competition vs. colonization tradeoff at intraspecific level. Mean seed mass estimated from seeds (n = 10-30 air-dried seeds) produced by 4-5 individuals per *A. thaliana* accession. Fecundity measured as the total number of fruits per accession multiplied by the average fruit length (n = 8 individual plants per accession). Each point represents the mean trait value of one *A. thaliana* accession (n = 71). Pearson’s correlation coefficient (*r*) and statistical significance (*P*) are shown, as well as the linear regression line (red).

### The competition vs. colonization trade-off shapes the center-to-margins differentiation of *A. thaliana* populations in Europe

The geographical groups of *A. thaliana* were phenotypically differentiated along its European distribution range. Specifically, we observed that plants from the Center nonrelict groups had 3-fold higher fecundities and were on average 12 and 6 cm taller than South relict and High-North nonrelict accessions, respectively (Fig. 3, Table 1). In contrast, accessions from the Center nonrelict groups significantly produced lighter seeds and their fecundity was more strongly reduced under biotic and abiotic stress conditions compared to groups from the range edges (up to 3-fold reduction in competition and water stress; see Fig. 3, Table 1). In between, South nonrelict and Mid-North nonrelict accessions exhibited intermediate trait values between central and peripheral ones, except for seed mass values, which were similar to central groups (Fig. 3).

**Table 1.** Outputs of the linear mixed models testing differences in fecundity, plant height, seed mass, and reaction norms in terms of fecundity in response to water stress and intraspecific competition among *A. thaliana* accessions by geographic group. R^2^_m_: Squared-R for fixed factors and R^2^_c_: Squared-R including random factors. Data shown are the F-values, the degrees of freedom (df) and the statistical significance level of each model using Type II tests. * *P* < 0.05, ** *P* < 0.01, *** *P* < 0.001.

| | Geographical<br>group<br>*df*= 4 | $R^2_m$ | $R^2_c$ |
| --- | --- | --- | --- |
| *Fecundity*<br>(number of fruits x mean fruit lengths) | 55.81*** | 0.29 | 0.68 |
| *Max. plant height (cm)* | 23.95*** | 0.17 | 0.66 |
| *Seed mass (mg)* | 59.61*** | 0.36 | 0.65 |
| *Response to water stress* | 5.00 | 0.03 | 0.49 |
| *Response to competition* | 38.24*** | 0.29 | 0.86 |

**Fig. 3.**
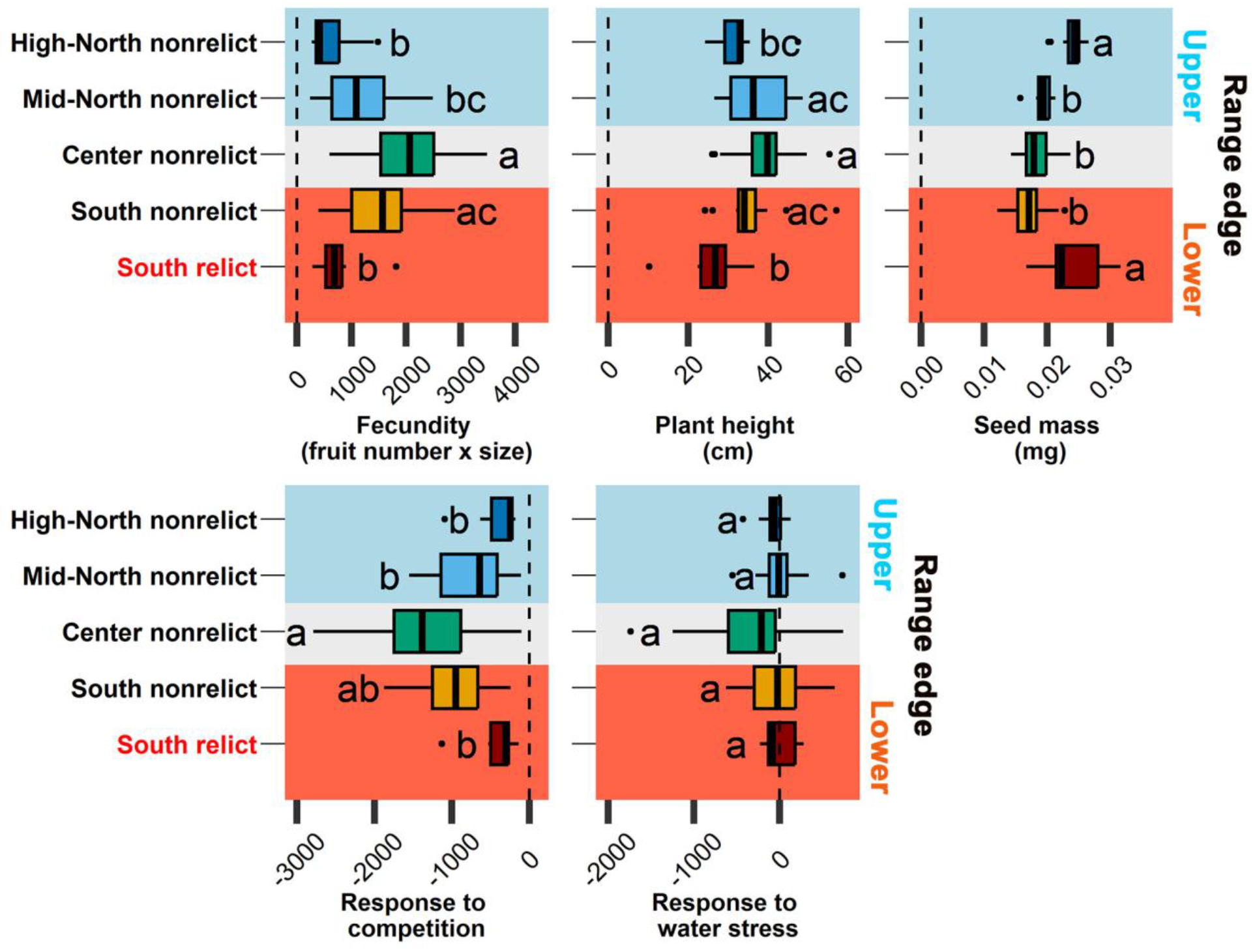
Phenotypic differentiation among *A. thaliana* geographical groups originating from central to range edge areas of its distribution range in Europe. Phenotypic differences in fecundity, plant height, seed mass and responses to intraspecific competition and water stress are depicted. Bars represent the average trait value (± SE) among accessions within each geographical group. Different letters indicate significant differences in the average trait among geographical groups after Tukey’s HSD test.

### Genetic convergence between edges populations at genes involved in dispersal and competitive ability

Using a polygenic GWAS approach^35^, we calculated the quantitative effect of 471,453 SNPs along the genome on each studied trait. SNP with positive effects on plant fecundity had also a positive effect on plant height (*r* = 0.04, *P* < 0.001; Table S2). In addition, SNP effects on fecundity were negatively correlated with SNP effects on seed mass (*r* = 0.04, *P* < 0.001; Table S2) and the responses to biotic and abiotic stresses (*r* = 0.03, *P* < 0.001 for competition response and *r* = -0.04, *P* < 0.001 for water stress response; Table S2). Overall, this showed that the phenotypic correlations observed between traits hold at the genetic level.

We estimated the quantitative genetic differentiation (*Q*_*ST*_) of traits among geographic groups for our traits and compared it to the distribution of allelic differentiation along the genome among geographic groups (*F*_*ST*_). The *Q*_*ST*_ values for seed mass, fecundity, plant height and the response to competition were substantially extreme in the distribution of *F*_*ST*_ across the genome (Fig. S4). It indicated that the structuration of genetic variance of these four traits among geographic groups could be associated to selective processes. We then extracted the 1% ^36^ (*i*.*e*. 4,715) SNPs equally distributed between the strongest positive and negative effects on every trait (*i*.*e*. 0.5% of the SNPs having the most negative effect, and 0.5% of the SNPs having the most positive effect). Using these sets of top-SNPs for every trait, we measured SNP-level *F*_*ST*_ between our geographical groups to estimate trait *F*_*STQ*_. In addition, we measured *F*_*ST*_ of non-coding SNPs to estimate *F*_*ST*_ at neutral markers. The *F*_*STQ*_ / *F*_*ST*_ ratio, i.e. the divergence of quantitative traits from neutral molecular markers at genomic level, revealed that seed mass has positive values higher than 1 (Fig. 4A), which suggested that seed mass is under directional selection. Moreover, pairwise comparisons of *F*_*STQ*_ / *F*_*ST*_ ratio between geographical groups showed that South relict group was genetically (at genes related to seed mass) closer to High-North nonrelict, and farther from Center nonrelict accessions. Similar although weaker patterns were also observed for fecundity and plant height (Fig. 4A). On the contrary, the *F*_*STQ*_ / *F*_*ST*_ ratio for biotic and abiotic stress responses was below 1 (Fig. 4A), suggesting that these traits are under uniform selection across the geographic range of *A. thaliana*^33^.

**Fig. 4.**
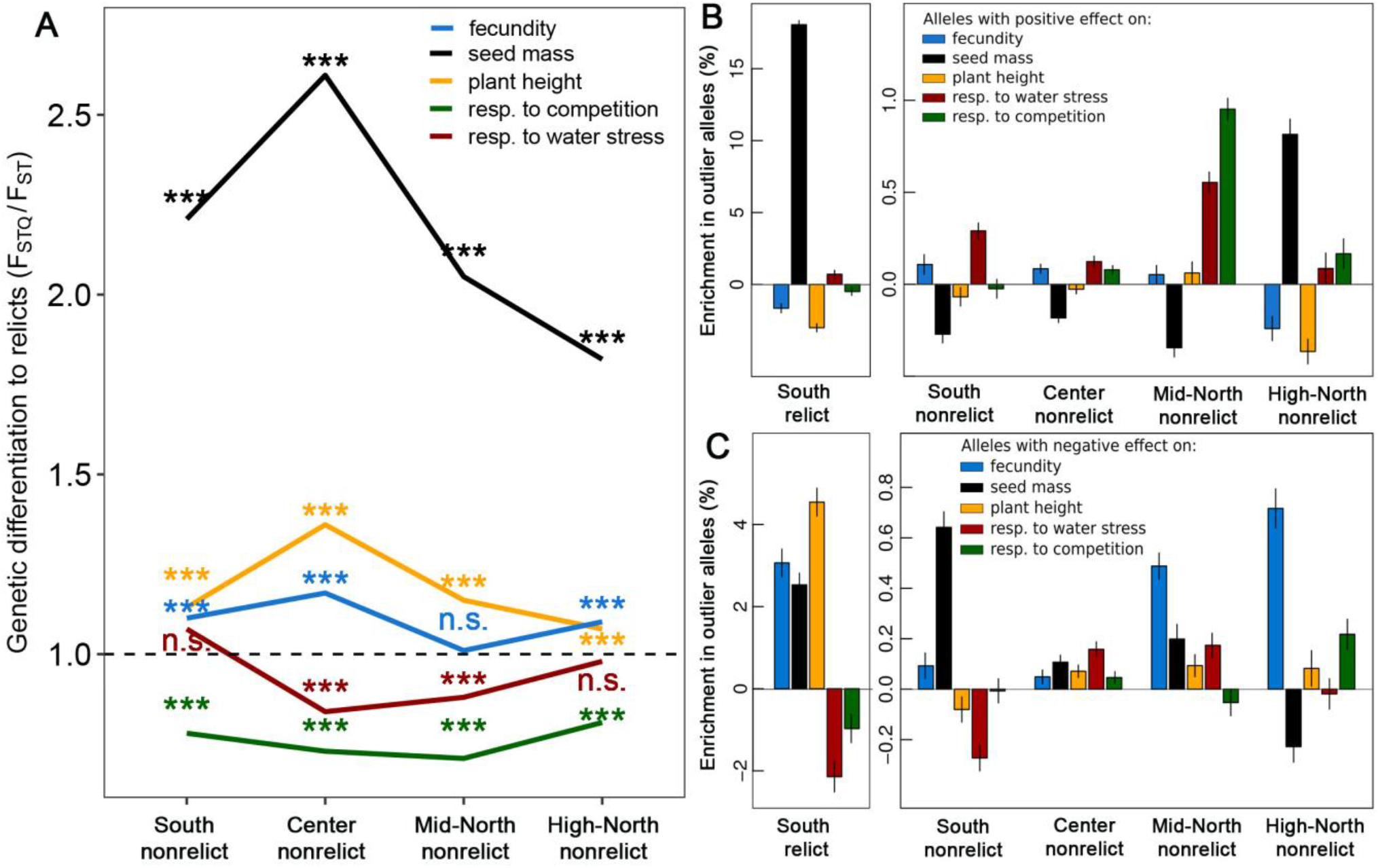
Genetic differentiation between genotypes originating from central and range edge areas. (A) Pairwise *F*_*STQ*_ / *F*_*ST*_ ratio comparisons between relict and nonrelict groups for fecundity, plant height, seed mass, response to water stress and intraspecific competition. The significance of *F*_*STQ*_ / *F*_*ST*_ ratio was tested with a simple linear model through comparing *F*_*ST*_ in non-coding regions versus *F*_*ST*_ in top-SNPs of a trait. n.s.: not significant; #: P < 0.1; *: *P* < 0.05; **: *P* < 0.01; ***: *P* < 0.001. (B and C) Enrichment in outlier haplotypes (%) with a positive (B) and negative (C) effect on life-history traits related to colonization (fecundity), competition (seed mass), dispersal (plant height), and response (resp.) to water stress and intraspecific competition for each geographical group. The “enrichment in outlier haplotypes (%)” for every trait and geographical group was calculated as the ratio between the proportion in outlier haplotypes in top-SNPs and those in non-coding regions.

Using published data from Lee and colleagues^24^, we then examined whether the 1% top-SNPs involved in the different traits measured were enriched or impoverished (compared to a random subset of SNPs) in outlier haplotypes in populations that are close to the range edges. As expected, there was a strong enrichment of outlier haplotypes in SNPs having a positive effect on seed mass for South relict and High-North nonrelict genotypes, while these SNPs were depleted in outlier haplotypes in Central nonrelict populations (Fig. 4B). Inversely, the SNPs having a negative effect on seed mass were significantly depleted for outlier haplotypes in High-North nonrelict genotypes, while there were enriched in outlier haplotypes in central populations (Fig. 4C). Surprisingly, we observed that relicts had an enrichment of outlier haplotypes with negative effect on seed mass but it represented a percent substantially lower than the percent of enrichment of haplotypes with positive effect (2 vs. 18%, Fig. 4B and 4C), perhaps reflecting the high diversity within the relict group. Moreover, the percent of outlier haplotypes with negative effect decreased for fecundity and plant height and increased for stress response ability by more than ∼ 3% from center to peripheral areas (Fig 4C). Together, these results suggest that the genetic convergence between South relicts and High-North nonrelict groups is stronger at genes involved in dispersal ability such as seed mass and fecundity than in central nonrelict accessions.

## DISCUSSION

Trade-offs in major life-history traits are expected to impose limits to phenotypic evolution, which is critical for species range dynamics. Here we combined experimental phenotypic data with available genome-wide sequence data to elucidate the ecological and evolutionary drivers of the distribution of *A. thaliana* populations in Europe. Our study revealed that colonization ability was selected against competition and stress tolerance abilities, leading to population differentiation along the *A. thaliana* distribution range.

The current demographic distribution of *A. thaliana* in Europe is the output of two consecutive expansion waves after the last glaciation^24,25,37^. The first expansion was driven by relict populations expanding northwards from their glacial refugia located at southern areas. A second one (∼10 ky ago) was driven by a nonrelict group that recolonized from Central-Eastern Europe to the South and North of Europe. This nonrelict group mainly erased any trace of relict groups in mid latitudes, becoming cosmopolitan in Europe. Previous studies speculated that such spatial genomic distribution was made possible by advantageous phenotypes allowing to nonrelict groups to carry out species range expansion and to relict groups to persist in the southern geographic limits^24,37–39^. Our study corroborates that the competition-colonization trade-off imposed phenotypic limits to *A. thaliana* populations across its geographical range (Fig. S5). One end of the trade-off is represented by the cosmopolitan group (Central nonrelict group), which exhibited a high colonization ability allowed by their taller stems and their production of smaller but more numerous seeds in comparison to relict groups. At the other end of the trade-off are populations occupying range edges (relicts from the South and High-North nonrelict group), which presented a high tolerance to competition associated with the production of big few seeds and short stature. Consistently, Estarague and colleagues^40^ recently showed phenotypic differentiation in leaf traits between central and peripheral *A. thaliana* populations across Europe. Moreover, Clauw and collaborators^41^ showed that northern peripheral accessions took advantage of their big seeds to deal with cold temperatures and survive winter stress. In addition, these authors showed that big seeds in northern populations were associated with lower growth rate, which is consistent with other findings in which slow growing and long-lived plants were preferentially found at the edges of the geographical range, while fast growing and short-lived ones were more abundant in central habitats^42,43^.

As a plant model species, *A. thaliana* offers a wealth of molecular background information to delve into evolutionary mechanisms that shape phenotypic relationships. First, our polygenic GWAS analysis evidenced that SNPs positively linked to colonization function had a pleiotropic and negative effect on competition and stress response abilities (see SNPs correlations on the involved traits; Table S2)^44^. In other words, this suggests that both colonization and competition functions could not be selected for in the same genetic background (but see ^45^ with recombinant inbred lines of *A. thaliana*). Second, *F*_*STQ*_ / *F*_*ST*_ comparisons among geographical groups revealed that selection differently favors seed mass, plant height and fecundity at the center and at the range edges, highlighting seed mass as key adaptive component to survive and persist in peripheral areas^41,45–47^. More precisely, we detected an enrichment of outlier haplotypes, i.e. an increase in outlier haplotypes compared to a random subset of SNPs along the genome, with positive effect on seed mass and intraspecific competition tolerance in South relict and High-North nonrelict accessions. By contrast, the same geographic groups were depleted in outlier haplotypes positively linked to dispersal and colonization ability. Moreover, a depletion of outlier haplotypes with negative effect on response to water deficit was detected from central to peripheral areas, in concordance to findings by Exposito-Alonso and collaborators^38^. In between, South and Mid-North nonrelict groups showed a higher enrichment in outlier haplotypes with positive effect on biotic and abiotic stress tolerance in comparison to central nonrelict group, but they decreased in outlier haplotypes positively linked to seed mass. These results suggest a potential introgression of outlier haplotypes from groups with high amount of outlier haplotypes (relicts and High-North group) to nonrelict groups in areas where both coexist (e.g. Iberian Peninsula and Sweden). This may have allowed more recent colonizers to obtain locally adaptive alleles for surviving and persisting in areas where the environmental limits of the species arise^27,38,45^. Evidence of genetic admixture between relict and nonrelict groups have been found in genes linked to flowering traits^48^. Our study lead us to speculate that genetic admixture in major life-history traits may have promoted the recolonization of *A. thaliana* throughout of the entire continent^24,37,49^.

In a broader sense beyond an evolutionary and ecological context of *A. thaliana*, our study provides unique and compelling evidence about the colonization-competition tradeoff as a mechanism structuring phenotypic diversification at intraspecific level, which has been overlooked in the literature^7,50,51^. This study confirms that divergent environmental selection across distribution ranges favors differential populations’ capacities for colonization and for competition and their associated plant’ functions (dispersal, stress-tolerance capacities). All in all, the evolutionary potential of ecotypes in colonization and competition abilities across distribution range is key to avoid population declines and local extinctions as well as to favor niche shifts to deal with changing environmental conditions^3,5,52^. But also, the evidence of genetic introgression and the ability of ecotypes to express differential adaptive phenotypes related to major plant’ abilities may alter the ecological networks of species interactions, which ultimately impact on community assembly, species coexistence and biodiversity patterns^53,54^.

## CONCLUSION

Our study demonstrates that the colonizing success of central populations of *A. thaliana* after the last glaciation can be explained by a high fecundity and dispersal ability (tall stems and light seeds). We also show that a high colonization ability is genetically associated with a negative effect on plant tolerance to competition and abiotic stress. However, natural selection has favored the maintenance of alternative ecological strategies at the opposite edges of the European distribution range: genotypes with low dispersal ability but high tolerance to water stress and competition were naturally selected. In between, the potential admixture groups resulting from the incorporation of relict alleles can explain the intermediate phenotypes between cosmopolitan (nonrelict) and relict accessions. Together, our results explain the center-to-margins genetic and phenotypic differentiation of *A. thaliana* populations. They also demonstrate the role of seed mass for plant adaptation across geographic gradients. More broadly, our study empirically confirms a long-standing hypothesis in ecology: the seed mass/seed number negative relationship reflects an evolutionary trade-off between competitive ability and colonizing ability.

## MATERIAL & METHODS

### Plant material

We initially selected a total of 72 natural accessions of *A. thaliana* from three geographical origin, namely central, northern and southern areas in Europe. Within each geographical group, we selected 24 natural lines that maximize the West-East axis distribution, with an altitudinal distribution below 1000 m a.s.l. to avoid confounding factors associated to elevation (Table S1). Within each geographical group, we separated accessions with relict origin or high relict genome introgression (hereafter called ‘South relict’ and ‘High-North nonrelict’ groups, respectively) from accessions of the nonrelict, cosmopolitan group depending on their geographic origin (hereafter South nonrelict, Center nonrelict and Mid-North nonrelict^24,29^ Fig. 1). All accessions were included in the initial germplasm of the 1,001 Genomes project (http://1001genomes.org/^29^), and seeds were supplied from the Eurasian Arabidopsis Stock Centre (NASC) and the Arabidopsis Biological Resource Center (ABRC).

### Greenhouse experiment setup and treatments

The experiment was carried out in two adjacent compartments of a greenhouse, where plants grew in individual pots (7 × 7 × 6.5 cm) filled with a 1:1 mixture of a commercial peat moss (Neuhaus N2) and vermiculite (medium grain 3-6 mm). We kept the plants in cold temperature (∼10 ºC) under well-watered conditions to ensure vernalization in all accessions (i.e. breaking the genetic suppression of flowering) for the first 40 days (from end of January to mid-March 2020). After the vernalization period, we raised the room temperature (20 ºC day / 15 °C night), and equally divided them into three different treatments differing in intraspecific competition and soil moisture conditions. Such environmental conditions have been widely proved to vary across biogeographical gradients^55^ and to exert selection for different *A. thaliana* traits and plastic responses^30,56– 58^. In the control treatment, a single individual was grown per pot under well-watered – regular irrigation every 4-5 days along the experiment – and no competition. In the intraspecific competition treatment, a focal *A. thaliana* plant was grown at the pot center surrounded by four individuals of the same accession, under similar well-watered conditions as the control treatment. In the water stress treatment, a single individual was grown in a pot, i.e. without competing neighbors, and in water stress conditions - regular irrigation every 10-11 days along the experiment since plants reached three - four true leaves (Fig. S1). Plants were harvested once they reached maturity, i.e. when the older fruit was senescent (from mid-April to mid-August 2020).

We used eight replicates of each accession and treatment, resulting in a total of 72 accessions x 8 replicates x 3 environments = 1,728 pots. Plants were equally distributed in the two adjacent compartments, with three large tables each and containing one treatment per table and compartment. Each table was divided into four similar squares, that included one replicate per accession randomly placed. Moreover, the tables were rotated within the compartments and turned around themselves every two days to minimize putative microclimate heterogeneity in each compartment.

### Phenotypic characterization of A. thaliana accessions

For each focal individual, we measured the maximum reproductive height (cm) from the rosette base to the apex of the longest flowering stem^10^. We counted the total number of fruits produced for each focal individual and measured the average fruit length from four fruits chosen randomly along the main inflorescence. We then calculated plant fecundity by multiplying the total number of fruits by the average fruit length^59^. On the other hand, we measured seed mass (BALCO MC5, mg) of focal individuals (n = 4-5 per accession) by weighing air-dried seeds (n = 10-30) and then dividing the total air-dried weight by the number of seeds in the sample. Finally, we assessed the stress response of each focal individual through the reaction norms of fecundity as the difference in fecundity of a particular individual in each stressful environment (intraspecific competition or water stress) to the mean fecundity of all focal individuals per accession in control conditions.

### Statistics

#### Phenotypic analyses

From the initial 1,728 individuals, we finally conducted analyses over a total of 1,600 individuals, after discarding one of the selected natural accessions from the Center nonrelict group due to absence of germination (Table S1) and focal plants that did not complete their life cycle at the end of experiment or died during the experiment. In total, 545 plants grew under control, 508 grew under intraspecific competition, and 547 were under water stress conditions. In the competition treatment, we did not consider in our analyses the focal plants with absence of one or more of their four neighbors to avoid potential bias linked to neighboring plant density. Further, we removed 22 individuals that showed extreme values within each accession and environment through applying the Hampel filter, i.e. the median, plus or minus 3 median absolute deviations^60^.

We first describe phenotypic correlations between our studied functional traits (plant height, fecundity, seed mass and stress responses). To assess differences in plant dispersal, colonization and competition abilities among *A. thaliana* geographical groups, we ran linear mixed-effects models for plant height, fecundity, seed mass on individuals in control conditions. We used the geographical group as fixed factor and accession identity (nested within geographical group) and squares (nested in table and these, in turn, nested in compartment) as random factors. With the same fixed and random structure, we run linear models for the reaction norms to both intraspecific competition and water stress among geographical groups. When the fixed factors turned out significant, we assessed mean phenotypic differences among geographical groups through *post hoc* Tukey’s tests. All analyses were performed in R v. 3.5.1 ^61^, using lme4 ^62^ and emmeans ^63^ R packages.

### Genetic analyses

We downloaded sequenced data from the 1,001 Genomes Project and filtered SNPs with a minor allele frequencies (MAF) inferior to 5% among the 71 accessions (∼ 1.2 million SNPs after filtering). With individual PLINK BED files containing genomic and phenotypic information for each studied trait, we run independent Bayesian sparse linear mixed models (BSLMM) implemented in the software package GEMMA^35^. BSLMM is a polygenic model that assesses the contribution of multiple SNPs to phenotypic variation, accounting for relatedness via inclusion of a kinship matrix as a covariate. From the BSLMMs, we obtained a data set with 471,453 SNPs, which was subsequently used to estimate join effect of these SNPs on each studied phenotypic trait. BSLMM models two effect hyperparameters: a basal effect α_i_, which captures the fact that many SNPs contribute to the phenotype, and an extra effect, β_i_, which captures the fact that a subset of SNPs has stronger effects than the rest. We summed α_i_ and β_i_ to estimate the effects of all SNPs. Finally, we performed linear correlations between all pairwise combinations of SNP effects on the phenotypic traits to assess whether ecological trade-offs were embedded at genetic level.

We tested the adaptive nature of phenotypic differentiation among geographic groups by comparing the structuration of genetic variance of traits among populations (*Q*_*ST*_) to their neutral genetic differentiation. To proceed, we firstly estimated Weir and Cockerham *F*_*ST* 64_ between each pair of geographic groups for all SNPs across the genome using PLINK software. *Q*_*ST*_ of each trait was then calculated as the ratio of genetic variance of each trait within and among geographic group. We tested if *Q*_*ST*_ values were extreme in the distribution of *F*_*ST*_ in a bootstrapped and non-parametric approach. In parallel, we extract, from BSLMMs, the 0.5% SNPs with the strongest positive effect and the 0.5% SNPs with the strongest negative effect on every trait, making a total of the 1% top-SNPs (i.e. 4715 SNPs per study trait). Additionally, we extracted the SNPs falling in non-coding regions (gene positions were downloaded from arabidopsis.org). The *F*_STQ_ parameter was estimated for each trait as the mean *F*_*ST*_ of the 1% top-SNPs for the given trait. (*F*_*STQ*_ / *F*_*ST*_ test) ^33^. We estimated the significance of *F*_*STQ*_ / *F*_*ST*_ ratio with a Student’s t-test (*F*_ST_ in non-coding regions versus *F*_ST_ in top-effect-SNPs of a trait). Finally, we quantified the *F*_*STQ*_ */ F*_*ST*_ ratio differences between all nonrelict groups and the ‘South relict’ group for every trait.

### Enrichment of outlier haplotypes for each phenotypic trait and geographical group

Using the calculation of the genomic proportion of relictiveness (i.e. ancestry proportion to the relict group) of each accession previously quantified by Lee and collaborators ^24^, we estimated the proportion of outlier haplotypes among our selection of 1% SNPs with the strongest positive and negative effect on each studied trait and among SNPs in non-coding regions. To do so, we applied a bootstrap with 50 permutations, each taking 1,000 SNPs among the 0.5% positive effect and others 1,000 SNPs among the 0.5% negative effect on a trait, and 1,000 SNPs randomly chosen among non-coding SNPs (to represent the neutral genomic background). Then, we estimated the average proportion of outlier haplotypes for these three categories (top-positive, top-negative and non-coding) among all accessions within each geographical group. The “enrichment in outlier haplotypes (%)” for every trait and geographical group was calculated as the difference between the proportion in outlier haplotypes in top-SNPs and those in non-coding regions.

## ACKNOWLEDGMENTS

We are very grateful to Estelle Leca, Estela Illa Bachs, Stefania Przybylska for their help in the setting of the experiment. We also thank Anais Hany and Thibault Martino for their valuable help in measuring phenotypic traits. We thank the technical platform “Plateforme des Terrains d’Expérience du LabEx CeMEB” in Montpellier for their support and advices during the experiment. This work was supported by CNRS, INRAE, the French Agency for Research (ANR grant ANR-17-CE02-0018-01; “AraBreed” to FV, DV, and CV), and the European Research Council (ERC;“CONSTRAINTS”: grant ERC-StG-2014-639706-CONSTRAINTS to CV). CCB was supported by a postdoctoral fellowship of the Ramón Areces Foundation and CCB was also supported by a postdoctoral fellowship of the Junta de Andalucia (Spain) and the European Social Fun 2014-2020 Program (DOC_01035). M.E.-A. is supported by the Office of the Director of the National Institutes of Health’s Early Investigator Award with award number: 1DP5OD029506-01; by the U.S. Department of Energy, Office of Biological and Environmental Research, grant number: DE-SC0021286; and by the Carnegie Institution for Science.

## AUTHOR CONTRIBUTIONS

CCB, AE, DV, FV, CV conceived and designed the experiments. CCB and AE performed the common garden experiment. CCB, AE and DV carried out the lab measurements. CCB, FV, AE analyzed the data. C-RL provided genomic data about proportion of relictiveness of each accession and together with ME-A gave support with statistical analyses. CCB, AE and FV wrote the paper with comments from all authors.

